# Spatial organization of the somatosensory cortex revealed by cyclic smFISH

**DOI:** 10.1101/276097

**Authors:** Simone Codeluppi, Lars E. Borm, Amit Zeisel, Gioele La Manno, Josina A. van Lunteren, Camilla I. Svensson, Sten Linnarsson

## Abstract

The global efforts towards the creation of a molecular census of the brain using single-cell transcriptomics is generating a large catalog of molecularly defined cell types lacking spatial information. Thus, new methods are needed to map a large number of cell-specific markers simultaneously on large tissue areas. Here, we developed a cyclic single molecule fluorescence in situ hybridization methodology and defined the cellular organization of the somatosensory cortex using markers identified by single-cell transcriptomics.

---

Recent increase in single-cell RNA sequencing (scRNA-seq) throughput and sensitivity have stimulated efforts to unravel the molecular characteristics of the cell types that form the nervous system. Large-scale efforts are underway with the measurement of millions of single cell transcriptomes leading to the identification of large numbers of molecularly-defined cell types. However, even though scRNA-seq data can provide information on the cell type composition of a tissue, it lacks the spatial information required for the reconstruction of tissue architecture at a cellular level. Combinatorial sets of tens to hundreds of marker genes can be used to distinguish cell types, but methods are needed that can detect such marker sets in brain tissue with high sensitivity, accuracy and throughput.

Although methods to detect RNA and protein in situ have been a mainstay of biology for decades, classical methods such as in situ hybridization and immunofluorescence are only semiquantitative and do not afford high levels of multiplexing. By resolving individual mRNA molecules, it is possible to greatly improve sensitivity and quantitative accuracy, as well as to increase multiplexing^1^. Expressed RNA can be mapped in cells or tissue sections by in situ sequencing^2–5^, or by single molecule fluorescence in situ hybridization (smFISH)^6–8^. smFISH not only has the advantage of nearly 100% RNA detection efficiency^9–11^, but also can be multiplexed by both spatial and temporal barcoding^6,7,12^. However, marker gene selection in multiplexed smFISH methods is limited by either, optical crowding, where signal dots start to overlap, or transcript length, so that biologically relevant marker genes cannot always be targeted, therefore restricting the mapping of scRNA-seq defined cell types^6,8^. We decided to overcome these limitations by developing a non-barcoded and unamplified cyclic smFISH method (osmFISH) in which the number of targets scales linearly. Although this targeted approach limits the total number of genes that can be analysed, osmFISH has a wide dynamic range in detection of gene expression allowing large freedom in marker selection, benefiting the biological interpretation. In osmFISH each RNA molecule is visualized as a diffraction limited spot after the binding of 20 nt long fluorescently labelled DNA probes^10,11^. Multiple transcripts are targeted at each round of hybridization, separated by fluorescent color. After image acquisition, the detection probes are removed by formamide melting in preparation for the next round of hybridization (**Supplementary Fig. 1a**). Thus, the number of targets equals the number of fluorescence channels times the number of hybridization cycles (here, 3×13). Importantly, since no barcoding is used, each image can be fully analyzed separately, and highly expressed genes (where spots are difficult to resolve) do not affect the detection of lower-expressed genes. This provides the freedom to design optimal probe sets without strong constraints on expression levels.

**Figure 1.**
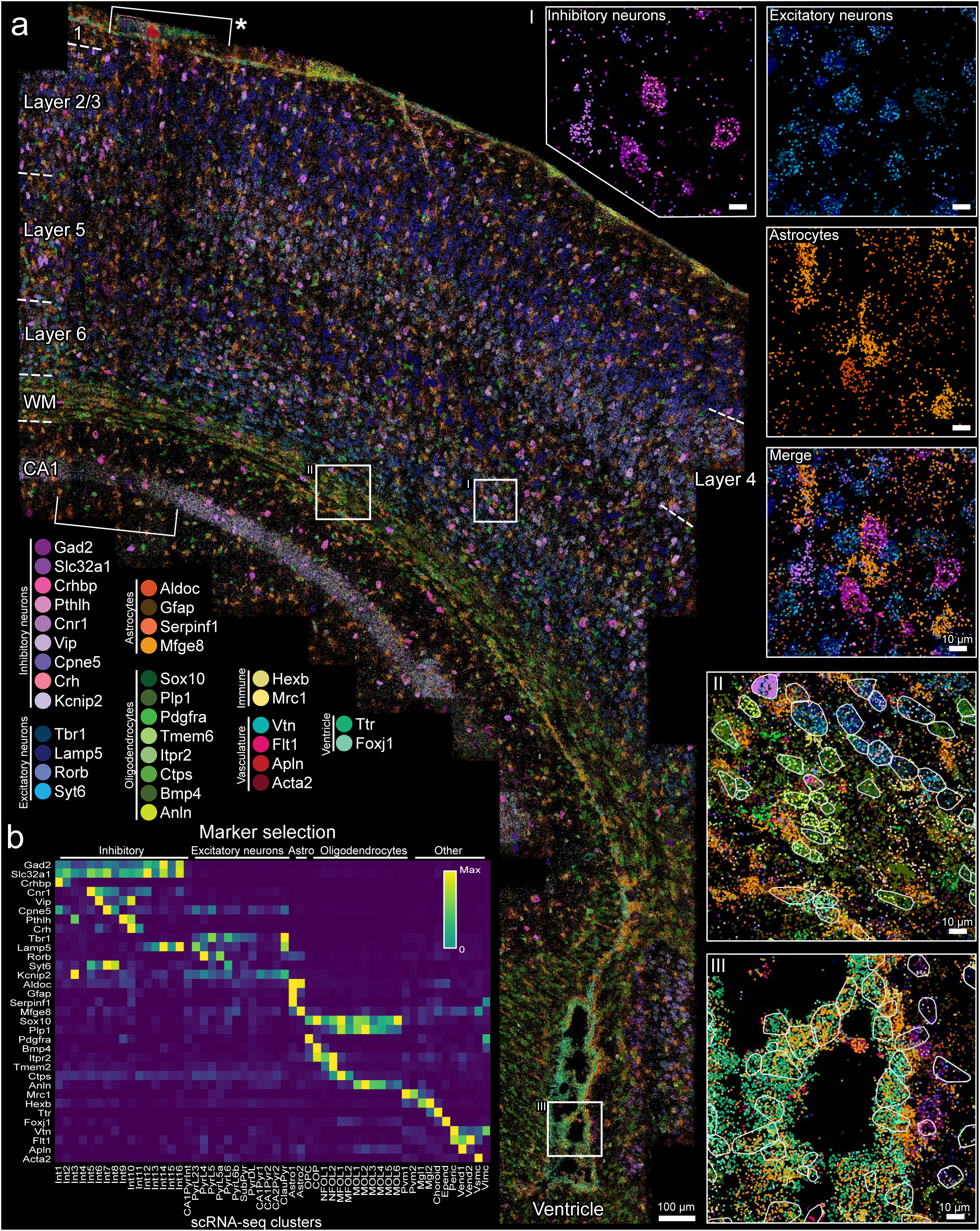
visualization of gene expression in the somatosensory cortex by osmFISH. **(a)** Image composite of 33 target genes. RNA molecules are visualized as dots of different colors according to gene. The combination of high-resolution imaging and large area coverage exposes both the structure of the tissue such as white matter (WM, II) and ventricle (III) and the fine RNA subcellular organization in different cell types (I). The white lines overlaid on the RNA images represent the segmented cells (II-III). The asterisk marks the region imaged after stripping and the dashed lines highlight the layers of the cortex, white matter and CA1 region of the hippocampus. **(b)** Heatmap of the markers gene expression level and the corresponding scRNA-seq clusters

osmFISH is characterized by a short hybridization time (2-4 hours), achieved through a heat shock to make the RNA more accessible, and high signal-to-noise ratio due to background reduction by tissue clearing. Furthermore, in order to maintain tissue integrity and RNA stability for experiments lasting multiple weeks, we implemented a method to covalently bind the brain sections to the coverslip. Additionally, to increase throughput and reduce human errors, we developed a semi-automated system in which the sample is positioned in a custom-made aluminium frame that fits up to 6 individual hybridization/imaging chambers and the reagents are dispensed by a computer-controlled fluidic system (**Supplementary Fig. 1b-e**) resulting in a staining protocol of 8 hours with minimal hands-on time. Finally, the key challenge for the application of smFISH based transcriptomics to biological questions, is processing the large, multi-terabyte, image datasets to obtain spatial single-cell gene expression profiles. Here we present a complete image analysis pipeline that we developed to process these large datasets, that is fully automated, efficient and easy to use, enabling the systematic analysis of spatial patterns at all scales.

As a first application, we used osmFISH to build an atlas of the mouse somatosensory cortex. To determine the identity of the cells in the tissue section, we selected 33 marker genes from a previously published scRNA-seq dataset of mouse somatosensory cortex^13,14^ (**Fig. 1b, Supplementary Table 1**). The markers were selected according to specificity and expression level in order to map all major cell types and most subtypes in 13 osmFISH cycles (**Supplementary Table 2**). A 3.8 mm2 tissue region including part of mouse somatosensory cortex, hippocampus and ventricle (**Supplementary Fig. 2**) was imaged in five fluorescence channels with 220 fields of view and 43-level Z stacks using an 100X objective to maximize the optical space. Imaging was the time limiting step, lasting around 14 hours per cycle. By optimizing the wash cycle, we achieved satisfactory stability of the RNA over the two-week long experiment (5.3±6.9% RNA molecules lost/cycle, **Supplementary Fig. 3,4**).

**Figure 2.**
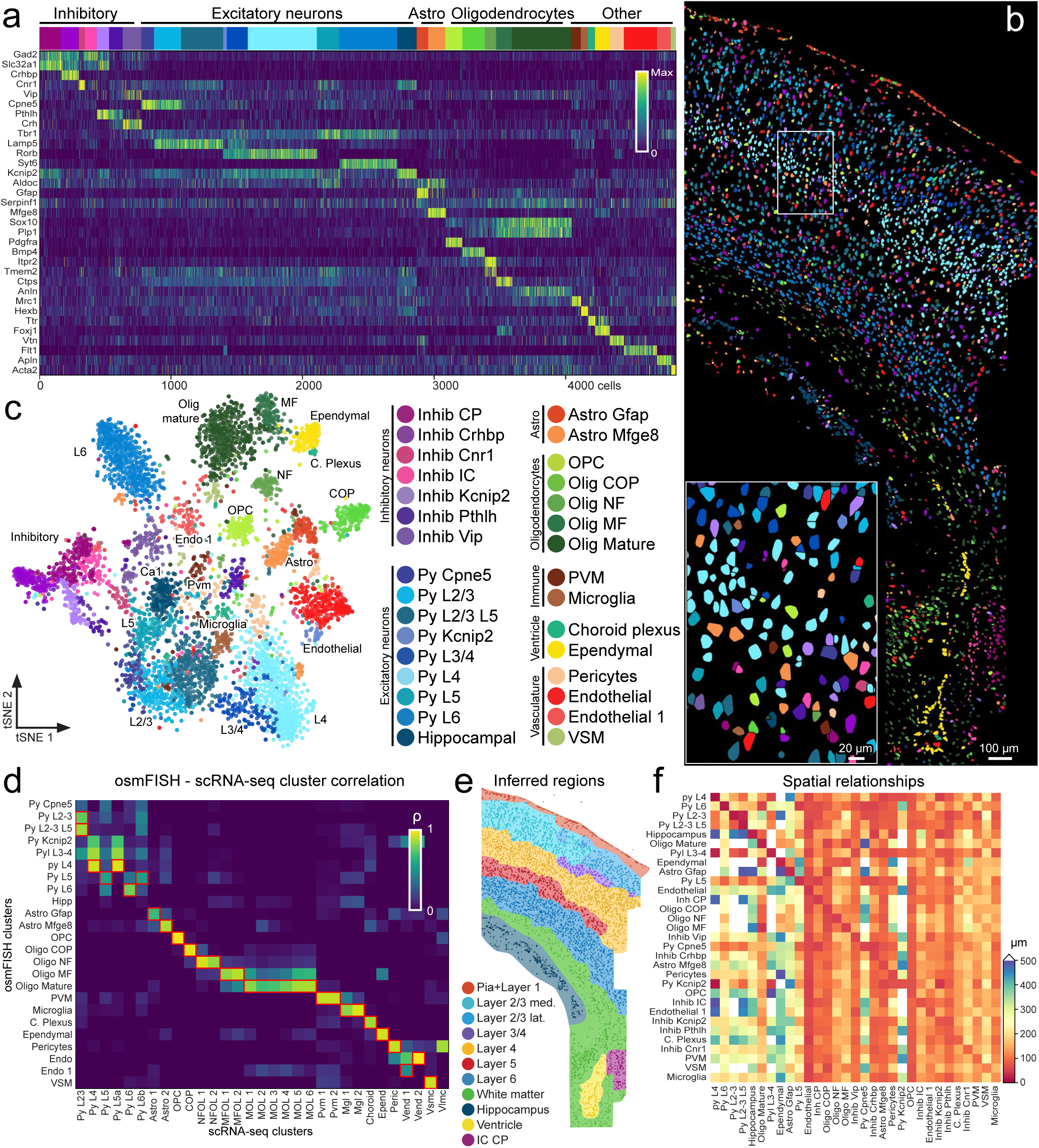
Clustering of osmFISH counts and mapping of the cell types. (a, c) Heatmap and t-SNE plot visualization of the 31 identified clusters with distinct expression profiles. (b) Spatial organization of the cell types in the tissue section as shown by coloring each cell according to the cluster. (d) Heatmap showing the correlation of the expression signature of corresponding osmFISH and scRNA-seq clusters, with best matches outlined in red. (e) Expression-driven spatial map of the somatosensory cortex. (f) Mean nearest neighbour distance between cells of all types.

A total of 1,976,659 RNA molecules were identified using the highly parallelised custom open source analysis pipeline (**Fig. 1a, Supplementary Fig 5**). The analysis pipeline ingests raw images, automatically stitches them, aligns the hybridization rounds and locates the RNA molecules. Cells were identified by segmentation of total mRNA visualized by a probe targeting poly(A) tails (**Supplementary Fig 6**). Single-cell expression profiles were obtained by counting RNA molecules in segmented cell bodies. The output is a matrix of counts similar to scRNA-seq data with the addition of the spatial coordinates of each cell and every RNA molecule mapped in the tissue section. The resulting dataset comprised 4839 cells. Molecule counts across cells were less sparse than the corresponding scRNA-seq data, showing only 30% zeros compared to 82% for scRNA-seq data^13^. Although the absolute number of zero measurements will depend strongly on the specific gene set, the comparative result indicates that osmFISH is substantially more sensitive than scRNA-seq. This was confirmed by the fact that osmFISH detected an average of 10.2±10.9 times more molecules per cell for the same set of genes analyzed^13^ (**Supplementary Fig. 3b**).

Spatially resolved data (e.g. osmFISH) and scRNA-seq data can be aligned either by probabilistic cell type calling using the scRNA-seq clusters as ground-truth reference, or by separately clustering the osmFISH expression profiles and then aligning the resulting clusters with those obtained by scRNA-seq. Because of the higher sensitivity and low dropouts of smFISH compared to scRNA-seq we choose to independently cluster osmFISH data in order to allow for de-novo discovery of cell types. We used a custom iterative clustering method (see Methods) to identify a total of 31 distinct clusters, most of which correlated clearly with a counterpart in the scRNA-seq defined cell types (**Fig 2 A-D, Supplementary Fig. 3c**). We observed astrocyte subtypes, microglia, choroid plexus epithelial cells, ependymal, pericytes, perivascular macrophages (PVM), vascular smooth muscle (VSM) and endothelial cells, as well as all types that form the oligodendrocyte lineage, including oligodendrocyte precursor cells (OPC), differentiation-committed oligodendrocyte precursors (COP), newly formed (NF), myelin forming (MF) and mature oligodendrocytes^13,14^ (**Fig. 2a,c**).

Neurons constitute 59% of our dataset and were subdivided into seven inhibitory (16%) and nine excitatory types (43%). The excitatory neurons followed the expected spatial positioning in the layered structure of the cortex, confirming the robustness of our clustering approach (**Fig. 2a,c, Supplementary Fig. 8**). Noteworthy, we were able to identify layer 5 excitatory neurons without including a specific marker in the osmFISH but using the expression of the pan-excitatory marker Tbr1 and the absence of layer specific markers together with their anatomical location. However, two types of excitatory neurons showed novel characteristics. First, even though the cells of the Py L2/3 L5 cluster were identified by a high level of expression of Lamp5 in layers 2/3, they were also found in layer 5, suggesting a multi-layer specificity for this cell type, or that our probe set could not fully detect the heterogeneity in this population (**Supplementary Fig. 8aI**). Second, the cells of cluster Py L3/4, were identified by joint expression of layer 2/3 and layer 4 markers (Lamp5 and Rorb, respectively). Interestingly, the cells were positioned at the interface between layer 2/3 and layer 4 suggesting the presence of a transition region between the two layers (**Supplementary Fig. 8aII**).

Even though we were not able to fully resolve all the inhibitory neurons subclasses because of limited coverage of our selected markers, we found five known cortical subtypes characterized by the expression of a single gene or a combination of multiple markers (**Fig. 2a-c**). We also identified two additional inhibitory neuron types that were not present in the scRNA-seq data (Inhib IC and Inhib CP) and showed a poorly defined expression profile but a distinct spatial localization in the internal capsule (Inhib IC) and the caudoputamen (Inhib CP) (**Supplementary Fig. 8b**). This demonstrate the power of using the spatial information inherent to osmFISH in improving the interpretation of expression profiles.

The knowledge of the cell types and their exact position in the tissue can be used for automatic, unbiased and data-driven construction of tissue atlases. Classically, anatomical atlases are manually annotated based on cell morphology, cell densities, low-throughput immunohistochemistry and in situ hybridization. Becauseofthelowthroughputofthesetechniques, multiple consecutive sections need to be processed in order to chart the regions that form the tissue. Using an iterative graph-based algorithm that we developed to find anatomical regions by determining the spatially dominating cell types on a single section, we show that osmFISH can be used to automatically delineate the regions of the tissue (**Fig. 2e and Supplementary Fig. 9**). Furthermore, because the cell types were mapped on the same single tissue section, osmFISH data also revealed the spatial relationships between the cell types that define the tissue architecture. For example, measuring the average distance between cells of the same type showed high spatial self-affinity of ependymal cells and region-specific cell types, and spatial self-avoidance of inhibitory neurons, microglia and astrocytes (**Fig. 2f** diagonal). Comparing instead cells of different types, we found strikingly that endothelial cells (i.e. blood vessels) were located in close proximity (64.7±9.5 μm) to all other cell types (**Fig. 2f)**. Thus, the ability to map the full complexity of the tissue in the same section enables the study of the spatial relationships between all cell types, that work in concert to give rise to tissue function.

While previously published protocols have focused on increasing the number of genes targeted using multiplexed smFISH techniques^6,8^, most commonly in cultured cells, here we focused on the complementary challenge of processing large tissue areas by reducing tissue background and building image-processing tools to handle large image datasets. Furthermore, with the goal of building cell type atlases, osmFISH offers high dynamic range of detectable gene expression levels, and freedom from the interference between genes commonly observed in barcoded methods. As a result, osmFISH permits more freedom in marker selection from scRNA-seq reference datasets facilitating cell type identification, and the resulting data can be used for systematic analysis of wide-area gene expression and cell type distribution. The ability to map the expression level of a large number of genes in a wide area of the same tissue section creates the opportunity for the development of data-driven reference atlases of healthy human tissue, but also for mapping alterations of tissue in diseases.

## METHODS

### Tissue collection

Animal handling and tissue harvesting followed the guidelines and recommendations of local animal protection legislation and were approved by the local committee for ethical experiments on laboratory animals (Stockholms Norra Djurförsöksetiska nämnd, Sweden, N 68/14). Postnatal day 22 wild type CD1 female mice were perfused with cold and oxygenated artificial spinal fluid solution. The brains were then harvested, snap frozen in Tissue-Tek O.C.T. (Sakura) and stored at -80°C until used.

### Probes design

For each of the selected markers we selected the transcript regions that showed reads in the single cell dataset and used them as a template to generate up to 48 probes using the online design tool from Biosearch technologies^1^. If the number of probes was too low, the mRNA reference sequences were selected from the UCSC genome browser (mouse GRCm38/mm10), aligned in order to determine the common sequence and added to the original template after removing the overlapping regions. The probes were directly labelled using Quasar 570, California Red 610 and Quasar 670 (Biosearchtech). Probe sequences are listed in Table 1.

### Coverslips functionalization

Coverslips (Marienfeld) were rinsed twice with distilled water for 20 min and incubated for 24 hours in concentrated nitric acid (Sigma Aldrich). The coverslips were then washed with distilled water 4 times for 1 hr, sterilized by autoclaving and stored in 95% ethanol. Before use, the coverslips were air dried and functionalized by incubation in 2% (3-aminopropyl) triethoxysilane (APES, Sigma Aldrich) in acetone for 1 min, followed by a 1 min rinse in water and air drying^2^. Functionalized coverslips can be stored in a dry environment up to a month.

### Tissue sample preparation

Thin tissue sections (10 μm) were cut on a cryostat and mounted on functionalized coverslips. Directly after mounting the sections were fixed for 10 min with PBS buffered 4% PFA (Merck), rinsed twice with PBS (Ambion), dehydrated by 3 min incubation in propan-2-ol (Sigma Aldrich), air dried and stored at -80°C until used.

### Imaging flow cell setup

To ensure repeated imaging of the same tissue area, the coverslip with tissue was mounted in an aluminium frame engineered to fit in the microscope stage. A 500μl flow cell was then assembled on the coverslip to which reagents can be dispensed by both pipetting or using a microfluidic pump (Dolomite Mitos P-Pump) connected to a computer and controlled by a custom script (Supplementary fig. 1b-e, availability see below). Except for the pump, the system was placed in an oven set at 38.5°C. The pump was connected to the flow cell with a tube running through a bottle of water positioned inside the oven in order to equilibrate the buffers to the oven temperature before entering the flow cell. The system was equipped with a separate line connected to a syringe, for manual purging of air bubbles and shut-off valves to close the fluid path when the flow cell was disconnected for imaging.

### osmFISH

After setting up the hybridization flow cell, the tissue section was rehydrated in 2X SSC (Sigma-Aldrich) for 5 min and cleared 2 times for 5 min with 4% SDS in 200mM boric-acid pH 8.5 (Merck). Tissue clearing was followed by 5 washes with 2X SSC for 1 min and 2 washes with TE pH 8 (Thermo). A heat shock was then performed for 10 min at 70°C followed by 3 washes with 2X SSC at room temperature. The tissue was then incubated for 5 minutes with hybridization mix (2X SSC, 10% w/v dextran sulfate (Sigma-Aldrich), 10% v/v formamide (Ambion), 1mg/ml E. coli tRNA (Roche), 2mM RVC (Sigma-Aldrich) and 0.2mg/ml Bovine Serum Albumin (Sigma-Aldrich)) at 38.5°C. Three different probe sets targeting 3 transcripts were added to the hybridization mix at a final concentration of 250nM (Stellaris, LGC Biosearch technologies). The probe containing hybridization mix was added to the flow cell and incubated for 4 hours at 38.5°C. The tissue was then washed 4 times 15 min with 20% formamide and 1mg/ml Hoechst (Sigma-Aldrich) in 2X SSC at 38.5°C followed by a single wash with 2X SSC. After injection of Slow Fade imaging medium (Thermo) the flow cell was transferred to the microscope for imaging (described below). After imaging the sample was then washed 5 times with 2X SSC and the probes were melted off their target by a 30 min incubation in 65% formamide in 2X SSC at 30°C followed by 5 washes with 2X SSC. To verify successful stripping, a small region of the tissue was imaged, which was later excluded of analysis (Fig 1a asterisk). In order to measure counting dropout or to cover the whole gene marker set the hybridization-stripping cycles were repeated 10 and 13 times respectively. After the osmFISH cycles the total mRNA was labeled using 500nM 30-nucleotide-long poly-T-Alexa488 (IDT) labelled probe in hybridization mix.

### Immunohistochemistry

The immunolabeling was performed inside the hybridization flow cell after the poly-A labeling was stripped off in 1.5 hour with the stripping buffer. The tissue was washed 5 times with TBS and blocked with 5% goat serum (Jackson immunoresearch) in 0.2% Tween (Merk)-TBS for 1 hr at room temperature. After blocking, the tissue was incubated overnight at 4°C with 1:100 Lectin-DyLight-488 (DL-1174 Vector) and 1:3000 rabbit-anti-GFAP (Z0334 Dako). After washing with 0.2% Tween-TBS, the tissue was incubated for 2 hrs at room temperature with 1:400 goat-anti-rabbit-Alexa647 (A-21245 Thermo) and then washed 3 times with 0.2% Tween-TBS, 2 times with TBS and then imaged after injection of slow fade (Thermo).

### Imaging

Imaging was performed on a standard epifluorescence microscope (Nikon Tieclipse, Nikon), equipped with amotorized stage (Nikon or HLD117 Prior), a Nikon CFI Plan Apo Lambda 100X oil immersion objective, a SOLA white light source (Lumencor) or a Spectra X 7-line LED light source (Lumencor) and Zyla 4.2 plus scMOS camera (Andor) for detection. The hybridization/imaging flow cell frame fit the stage with minimal displacement therefore the later displacement between imaging cycles was minimal and the same region could be image multiple rounds. For each cycle we imaged 220 xy positions with 10% overlapping and collected z-stack of 0.3 μm steps. The exposure times were 30ms for Hoechst, 200 ms for FITC and 1s for all remaining channels (Quasar 570, California red 610, Quasar 670). During imaging the hybridization chamber was cooled to approximately 15°C with cold air.

### Image processing

Image processing was performed using a custom Python pipeline, which is freely available as open source (see below). All analysis steps were parallelised due to the large size of the dataset and performed on an MPICH running cluster.

Illumination correction was performed for each channel in every hybridization. The illumination function was determined by calculating the max projection of the gene average image after filtering with a 3D gaussian with large standard deviation. Each xy 3D tile was then normalized with the illumination function and the background calculated by blurring the illumination-corrected image with a gaussian with standard deviation larger than the smFISH dots or the nuclei depending on the channel processed. The nuclei signal of each hybridization was used to calculate the similarity transformation matrix used to stitch all the xy positions together, using a modified version of Preibisch et al.^3^. For RNA counting, after background subtraction the dots were enhanced with a laplacian of a gaussian with sigma smaller than the dots. The RNA dots were defined as the local maxima above a threshold automatically calculated, after removal of connected components larger than dots. The threshold was defined as the point that belongs to the distribution of the total number of local maxima calculated for different thresholds, with the maximum distance from the segment that connect the two extremities of the distribution. The raw RNA molecules in each hybridization were identified in the whole field of view of each xy tile and mapped to the stitched image using the previously calculated similarity transform (raw counts). After mapping the molecules for all xy tiles the duplicated dots in the xy overlapping regions were removed. After processing of all hybridizations, we used the stitched nuclei images to calculate a second similarity transformation matrix used to register all the hybridization. We then defined the cell objects by watershed-based segmentation of the total mRNA labelled in the last hybridization step. For each cell object the number and coordinates of RNA molecules for each gene was calculated as described above (cell mapped counts).

### Analysis of cell types

Cells included in the analysis were located outside the region imaged after stripping (Fig. 1a), had a size between 5μm2 and 275μm2 and contained at least one detected molecule. Data were normalized by total number of molecules of all genes per cell and the sum of each gene over all cells and subsequently multiplied by the total number of genes times cells. Cells were clustered with a custom iterative clustering algorithm using Scikit-learn agglomerative hierarchical clustering^4^. In brief, each iteration splits the dataset in two and evaluates whether the daughter-clusters are homogeneous (all cells having a similar expression profile) or heterogeneous (containing multiple homogeneous clusters). To decide if a daughter-cluster is heterogeneous, it is split into two granddaughter-clusters which are tested for differentially expressed genes. A differentially expressed gene is defined as having an expression level above 550 in at least one cluster (after normalization) and a two sided Mann-Whitney U test p-value below 0.1x10-20 comparing the enriched gene(s). Furthermore, a minimal cluster size of 15 cells was used. This process is repeated until all clusters are homogeneous. A split that separated a group of cells based only on an expression level difference was not allowed. However, not all cases were caught, so similar clusters were merged afterwards based on correlation.

To compare the osmFISH clusters with the clusters of the scRNA-seq, Pearson’s correlation coefficient was calculated for the mean expression levels of all cluster pairs. For visualization, t-SNE was used to embed the high dimensional data in 2D^5^.

### Spatial Analysis

To infer tissue regions based on the location of the cell types we first selected spatially organized clusters according to the Ripley’s K estimate^6^. Ripley’s K estimate was calculated on the centroids of all cells of each cluster with a radius of 750 pixels (∼50 μm) and the cluster with a Ripley’s K above 2x107 were selected. Then the cells in the sample were converged iteratively to the most common cluster label in their local neighbourhood, using a k=20 nearest neighbour network and a maximal distance that varied between 1000 and 1500 pixels (65-97.5μm) for 20 iterations. After iteration 5, 7, 10 and 16 a connected graph was made for each unique label with a maximum distance of 3000 pixels (195μm), and disconnected sub-graphs with a size smaller than 75 cells were reset to their original value, to prevent small local inflation of one label. Finally, isolated regions with the same label were split and cells that were far away from other cells were forced to adapt the identity of its closest 10 neighbours without maximum distance.

To visualize spatial relationships between cells of all types on the small scale, the distance of each cell to the closest neighbour of the same type (self-affinity) and all other types was measured. The mean distances of all permutations of cell types were plotted as a heatmap, where values above 500μm were excluded.

### Code

All code was written in Python and is available online: linnarssonlab.org/osmFISH The following Python packages were used: Astropy^7^, Dask-Distributed^8^, H5py^9^, Loompy, Matplotlib^10^, Mpi4py^11^, N2reader, Networkx^12^, Numpy^13^, Pandas^14^, Pyserial, Scikit-Image^15^, Scikit-Learn^4^, Scipy^16^ and Sympy^17^.

## Acknowledgments

We thank the Knut and Alice Wallenberg Foundation (2015.0041), the Swedish Foundation for Strategic Research (RIF 14-0057 and SB 16-0065), the Swedish Infrastructure for Computing (SNIC 2016/1-283), the Chan Zuckerberg Initiative (2017-174399) and the NIH (1U01MH114812-01) for generous support of this work. We thank Microliquid for help on design and manufacturing the flow cell. We thank Peter Lönneberg for maintenance of the computing cluster, Job van der Zwan for the loom-viewer visualization tool and Anna Johnsson for project management.

## Author contributions

S.C., L.E.B., S.L. designed the experiments, S.C., L.E.B., performed the experiments and data analysis. J.A.V.L. contributed to the image stitching code. G.L.M., A.Z. contributed to the analysis. L.E.B. designed the fluidic system.

A.Z. designed the hybridization chamber. C.I.S. contributed to funding acquisition and provided scientific feedback. S.C., L.E.B., S.L., wrote the manuscript. All authors provided critical feedback and helped shape the research, analysis and manuscript.

## Code and availability

All code and data is available online: linnarssonlab.org/ osmFISH. The raw osmFISH images, 6 million totalling to ∼5TB, are available upon request.

## Supplemental information

Supplemental information contains nine figures and two tabels.

